# Single-cell RNA sequencing identifies accumulation of Fcgr2b+ virtual memory like CD8 T cells with cytotoxic and inflammatory potential in aged mouse white adipose tissue

**DOI:** 10.1101/2025.06.27.661935

**Authors:** Archit Kumar, Martin O’Brien, Vincent B Young, Raymond Yung

**Author notes:** **Corresponding author:** Raymond Yung, Director, Geriatrics Center and Institute of Gerontology, Chief, Division of Geriatric and Palliative Medicine, Department of Internal Medicine, University of Michigan, Ann Arbor, USA-48109. **Co-corresponding author:** Archit Kumar, Division of Geriatric and Palliative Medicine, Department of Internal Medicine, Room no 3258, BSRB, 109 Zina Pitcher Pl, University of Michigan, Ann Arbor, USA-48109.

## Abstract

Aging and obesity are associated with pro-inflammatory changes in adipose tissue. Overlapping mechanisms, such as the infiltration of inflammatory macrophages and T cells into visceral adipose tissue, have been implicated in contributing inflammation. However, a comparative analysis from both states is needed to identify distinct regulatory targets. Here, we performed single-cell RNA sequencing of stromal vascular fractions (SVF) isolated from gonadal white adipose tissue (gWAT) of young mice fed either a normal or a high-fat diet, and aged mice fed a normal diet. Our analysis revealed that physiological aging, compared to high-fat diet induced obesity, was associated with accumulation of phenotypically distinct CD8 T cells resembling virtual memory (VM) CD8 T cells. These cells expressed high levels of *Cd44*, *Sell*, *Il7r*, *Il2rb*, lacked *Itga4*, and exhibited elevated *Fcgr2b* expression which was associated with pseudotime differentiation trajectories. Flow cytometry confirmed an age-associated increase in Fcgr2b+CD49d− VM like CD8 T cells in gWAT. Notably, these Fcgr2b-expressing cells exhibited a cytotoxic profile, and expressed granzyme M. Functional analysis using recombinant granzyme M revealed its potential in inducing inflammation in mouse fibroblasts and macrophages. Together, our study has identified Fcgr2b+CD49d− VM-like CD8 T cells in the adipose tissue of aged mice with regulatory, cytotoxic and inflammatory potential.

## 1 Introduction

Aging is an inevitable biological process characterized by the progressive deterioration of physiological functions, which can lead to increased vulnerability to disease and death. During aging, host immune cells lose their ability to respond to infections and cancer, while simultaneously acquiring a pro-inflammatory phenotype that contributes to tissue pathology. Moreover, chronic low-grade inflammation associated with aging, termed “inflammaging”, is a major driver of age-related diseases including type 2 diabetes, metabolic syndrome, cardiovascular diseases, and cancer (Leonardi et al. 2018; Franceschi et al. 2018; Ferrucci & Fabbri 2018). Many of these age-related diseases are also seen in obesity, suggesting that obesity can accelerate the aging process (Santos & Sinha 2021). Adipose tissue is one of the notable organs affected during aging, where age-related changes in immune cells are first detected (Schaum et al. 2020). Moreover, an increase in body weight, accumulation of visceral fat mass and adipose tissue dysfunction are associated with both aging and obesity (Ou et al. 2022; Reyes-Farias et al. 2021; Trim et al. 2018).

Adipose tissue (AT) is a metabolic active endocrine organ distributed throughout the body, playing a central role in regulating systemic metabolism through the secretion of adipokines and cytokines. AT comprises mainly brown adipose tissue (BAT) and white adipose tissue (WAT). BAT, predominantly located in the supraclavicular and interscapular regions, is essential for thermogenesis, while WAT functions primarily as an energy reservoir. The metabolic activity and plasticity of WAT is highly responsive to changes in energy supply and demand. WAT can be further subdivided into visceral adipose tissue (vWAT), which surrounds internal organs, whereas subcutaneous adipose tissue (scWAT), located beneath the skin. Accumulation of vWAT is strongly associated with increased risk of cardiovascular diseases, metabolic disorders and cancer (Medina-Urrutia et al. 2025; Zhang et al. 2025; Nguyen & Shanmugan 2024; Vasamsetti et al. 2023). Structurally, WAT is a complex and heterogeneous tissue composed of adipocytes, mesenchymal stromal cells, endothelial cells, and immune cells. Although adipocytes dominate tissue structural volume of WAT, a substantial portion of its cellular composition consists of non-adipocytes cells, collectively known as stromal vascular fraction (SVF). Functional changes in SVF of adipose tissue, especially fibroblasts and immune cells, play a critical role in regulating inflammation and metabolic homeostasis (Ou et al. 2022; Reyes-Farias et al. 2021; Trim et al. 2018).

Both aging and diet-induced obesity significantly alter the heterogeneity and functional of adipose tissue. These changes include secretion of senescent-associated secretory phenotype (SASP) proteins, acquisition of pro-inflammatory phenotype, accumulation of T cells, B cells and inflammatory macrophages in visceral adipose tissue (Camell et al. 2019; Lumeng et al. 2011; Winer et al. 2011; Nishimura et al. 2009; Weisberg et al. 2003). Recent advancements such as single-cell (sc) and single-nuclei (sn) RNA sequencing (RNAseq) have been employed to investigate these alterations in the vWAT, especially in gonadal white adipose tissue (gWAT), during aging and diet-induced obesity in mice (Wang et al. 2025; So et al. 2025; Wu et al. 2024; Liao et al. 2024; Kar et al. 2024; Muñoz-Rojas et al. 2024; Emont et al. 2023; Cottam et al. 2022; Sárvári et al. 2021; Mogilenko et al. 2021). Although recent studies have revealed important insights into the regulation of fibroblasts, regulatory T cells (Tregs) and exhausted T cells in adipose tissue, a comparative analysis of aging- and obesity-associated changes remains lacking. To address this gap, we conducted an integrated scRNAseq analysis of SVF isolated from gWAT of aged and obese mice of both sexes. Our findings revealed distinct differences in CD8 T cell memory subsets between aged and obese gWAT in mice. Specifically, memory CD8 T cells in aged mice express high levels of inhibitory receptor *Fcgr2b*, exhibits a cytotoxic profile, and could initiate granzyme M-mediate inflammatory responses in mouse fibroblasts and macrophages.

## 2 Results

### 2.1 Aging and high fat diet induce gWAT dysfunction in mice

Visceral adiposity induced during high fat diet (HFD) exposure or aging can influence the development of metabolic syndrome and systemic inflammation. To investigate the similarities and differences in adipose tissue dysfunction under these conditions, we assessed metabolic stress response and circulating adipokines levels in young mice subjected to normal diet (ND) or HFD (42% Kcal from fat) exposure for 12 weeks and aged mice maintained on ND. HFD exposure in young mice and physiological aging in ND-fed mice both significantly increased body weight and gonadal white adipose (gWAT) mass (Figure 1a, 1b, and 1c). Male mice generally exhibited significantly higher body weight compared to female mice. Notably, HFD-fed young male mice displayed greater adiposity, than their female counterparts, while aged female mice exhibited more pronounced fat accumulation (gWAT mass) than aged males. In response to metabolic stress, only HFD-fed young mice exhibited metabolic syndrome when compared to ND-fed young mice, as indicated by significantly elevated blood glucose levels during glucose tolerance test (GTT) and high fasting glucose levels (Figure 1d, S1a, S1b, and S1e). HFD-fed young male mice also showed increased serum glucose levels during insulin tolerance test, although the differences were not statistically significant when compared to ND-fed young male mice (Figure S1c, S1d, and S1f). Furthermore, male mice exhibited significantly worse outcome in during metabolic stress tests under both aging and HFD-induced obesity conditions, compared to females of the same cohorts.

**Figure 1.**
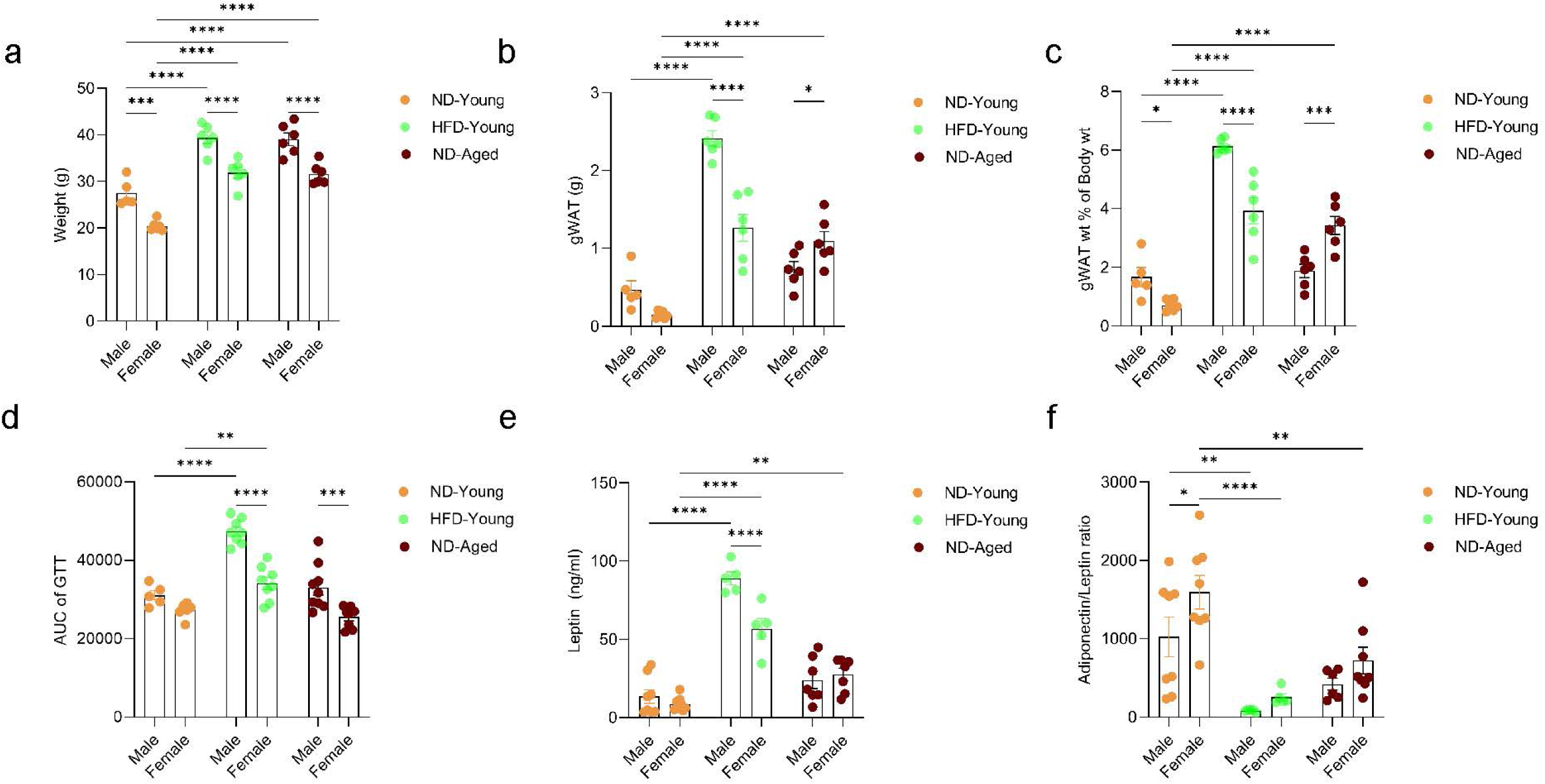
Aging and HFD-induced obesity impair gWAT function. (a) Body weight, (b) gWAT mass, (c) gWAT mass normalized to body weight in normal diet (ND)-fed young mice (4-5 months old), high fat diet (HFD)-fed young mice (42% Kcal from fat for 12 weeks, 4-5 months old) and ND-fed aged mice (21-24 months old) of both sexes. (d) Area under the curve (AUC) of serum glucose levels during a glucose tolerance test (GTT) following overnight fasting. (e) Serum leptin levels. (f) Serum adiponectin-to-leptin ratio. Data are presented as mean ± SEM. Statistical significance was determined using two-way ANOVA followed by Tukey’s HSD post hoc test. N = 5-8 mice per group. P-values < 0.05 were considered significant. p< 0.05 (*), p<0.01 (**), p<0.001 (***), p<0.0001 (****).

Dysregulation of adipokines levels, primarily adiponectin and leptin, are well established indicators of impaired adipose tissue function. To assess these changes under different physiological conditions, we measured serum adipokine levels in young mice subjected to ND or HFD and aged mice maintained on ND. HFD-fed young mice exhibited significantly elevated serum leptin levels and reduced adiponectin-to-leptin ratios relative to ND-fed young mice (Figure 1e, and 1f). Conversely, age-associated changes in the leptin levels were more pronounced in female mice compared to ND-fed young female mice. Female mice across all the experimental groups showed elevated adiponectin levels; however, no significant sex-specific differences in the adiponectin-to-leptin ratio were observed under either HFD or aging conditions, although only ND-fed young females exhibited significantly higher adiponectin-to-leptin ratio compared to ND-fed young mice (Figure 1e, 1f, and S1g). The results suggest sex specific regulation of adipokines and adiposity where female mice may exert more protective role against metabolic dysfunction than male mice. However, this protection appears to be diminished with aging and HFD-induced obesity.

### 2.2 Aged mice exhibit altered adipose tissue cellular heterogeneity

To characterize differences in adipose tissue heterogeneity during HFD exposure and aging, we performed scRNA-seq on the SVF isolated from digested gWAT of ND-fed young, HFD-fed young, and ND-fed aged mice of both sexes. SVF from each individual mouse (n = 3 per group per sex) was sorted to remove dead cells and equal numbers of viable cells from each mouse within a group were pooled to generate six distinct experimental groups (Figure 2a). Libraries were prepared using the 10x Genomics Chromium Single Cell 3′ platform and subjected to Illumina sequencing. After quality control and filtering, 32,806 cells were retained and integrated using Seurat. Dimensionality reduction with Uniform Manifold Approximation and Projection (UMAP) was used to visualize the data (Figure 2b). Cell type annotation was performed based on the expression of canonical and previously established marker genes (Figure 2c, and S2) (Liao et al. 2024; Muñoz-Rojas et al. 2024). Both HFD-fed young mice and ND-fed aged mice exhibited a reduction in adipose stem-like cells (ASCs) populations, particularly in males across all groups. ASCs heterogeneity has been shown to vary due to diet, age, sex, and plays a significant role in regulating adiposity (Wu et al. 2024; Liao et al. 2024; Kar et al. 2024; Sárvári et al. 2021). In line with previous literature, HFD-fed mice showed increased infiltration of macrophages whereas ND-fed aged mice demonstrated greater accumulation of B cells and CD8 T cells in gWAT (Camell 2022; Camell et al. 2019; Bénézech et al. 2015; Lumeng et al. 2011) (Figure 2d). The accumulation of CD8 T cells was more pronounced in ND-fed aged male mice, while B cell accumulation was higher in ND-fed aged female mice. Additionally, increased frequencies of γδ T cells, regulatory T cells (Tregs) were observed in ND-fed aged male mice (Figure 2d). These findings highlight distinct heterogeneity in the immune cells of adipose tissue during aging and diet-induce obesity.

**Figure 2.**
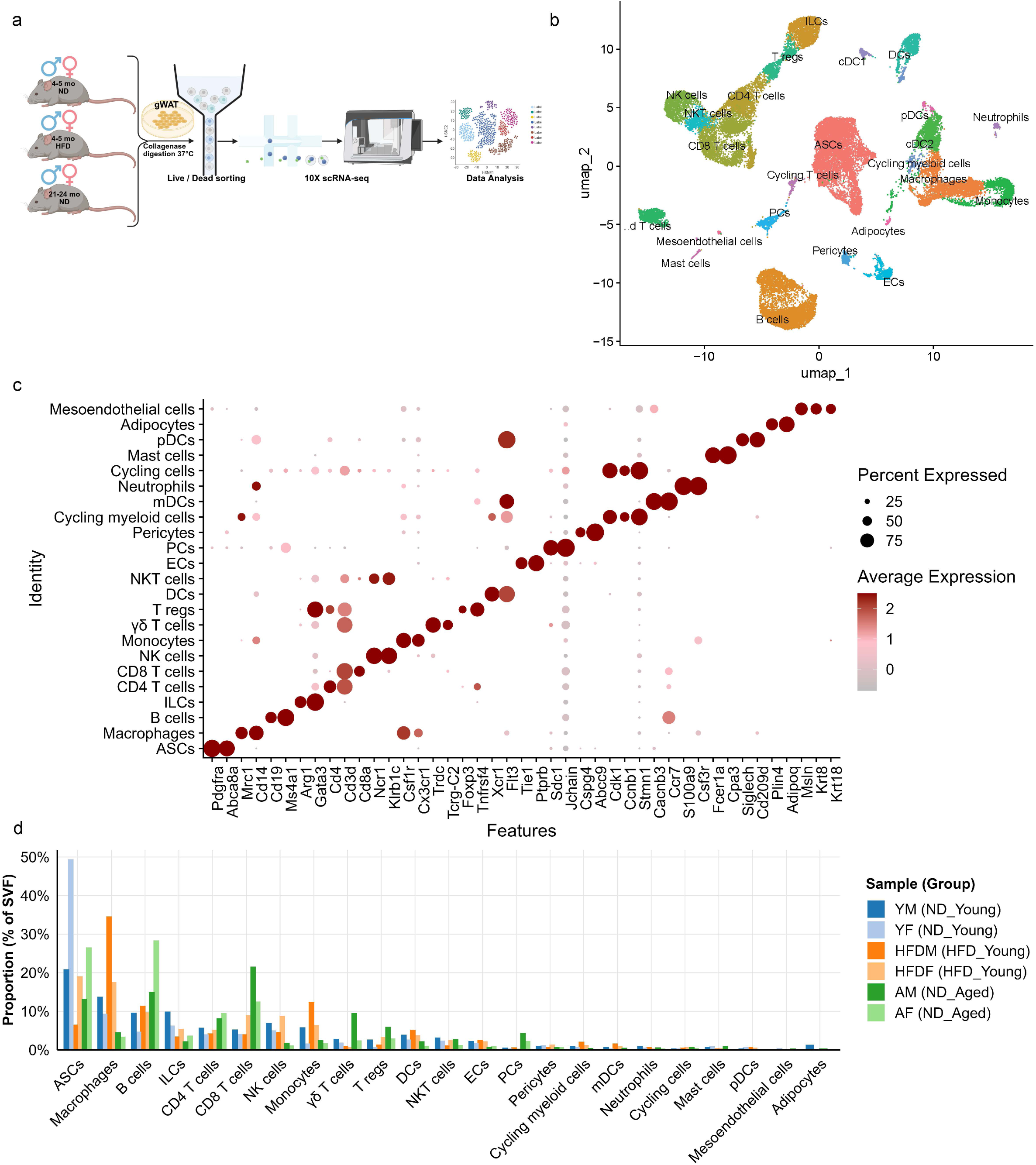
scRNA-seq analysis of stromal vascular fraction of gWAT. (a) Schematic overview of single-cell RNA sequencing (scRNA-seq) performed on SVF isolated from gWAT of ND-fed young, HFD-fed young, and ND-fed aged mice (both sexes), created using Biorender.com. Each scRNA-seq sample was generated by pooling SVF from three independent mice per group after live/dead cell sorting. (b) UMAP plot showing 32,806 cells after quality control and data integration, colored by annotated cell-types. (c) Dot plot showing expression of marker genes across identified cell-types. (d) Bar plot representing the proportions of non-immune and immune cell types in the SVF. ASCs: Adipose stem cells; ILCs: Innate lymphoid cells; NK: Natural killer cells; γδ T cells: Gamma delta T cells; T regs: T regulatory cells; DCs: Dendritic cells; NKT: Natural killer T cells; ECs: Endothelial cells; PCs: Plasma cells; mDCs: Migratory dendritic cells; pDCs: Plasmacytoid dendritic cells.

### 2.3 CD8 T cells in aged mice exhibit enhanced cell-cell communication

Following the identification of altered cellular heterogeneity in gWAT during aging and obesity, we next investigated cell-cell communication to identify interacting cell types. We employed CellChat, a computational tool that infers intercellular signaling networks based on curated ligand-receptor (L-R) interactions, to quantify signaling dynamics in young mice either fed ND or HFD and ND-fed aged mice (Jin et al. 2025). Our analysis revealed that CD8 T cells from both HFD-fed young mice and ND-fed aged mice displayed increased incoming signaling strength, indicating a higher degree of interaction with other gWAT cell types compared to ND-fed young mice (Figure 3a). However, no sex related changes were observed in the signaling strengths of cell-cell communication (Figure S3). Examination of incoming signals indicated that a broad engagement of CD8 T cells with various cell populations under both aging and HFD conditions (Figure 3b).

**Figure 3.**
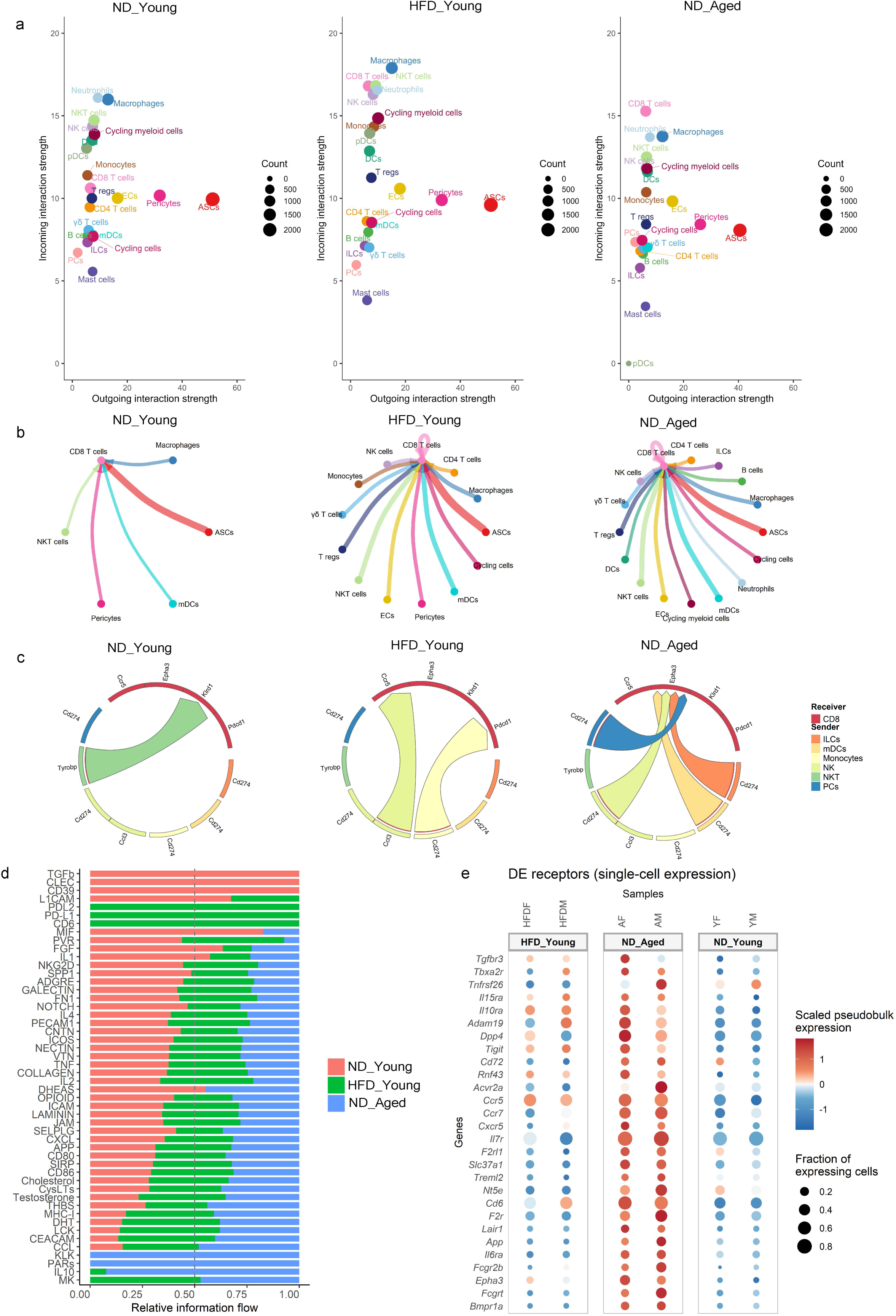
Cell-cell interaction within SVF. (a) Dot plots representing incoming and outgoing signaling strengths between cell types within the SVF of ND-fed young, HFD-fed young and ND-fed aged mice inferred using Cellchat. Dot size reflects the number of cells within cell type. (b) Circle plot showing incoming signals to CD8 T cells from other cell types based on ligand-receptor (L-R) expression. Each node represents a cell type, and edge width indicates interaction strength. (c) Chord diagram representing incoming signals to CD8 T cells inferred using MultiNicheNet. The top 100 differentially expressed L-R pairs among ND-fed young, HFD-fed young and ND-fed aged mice were used to generate the plots. Edge width represents the predicted strength of incoming signals to CD8 T cells. (d) Bar graphs showing the strength of signaling pathways targeting CD8 T cells in each group, generated using rankNet() function in Cellchat. (e) Dot plot displaying receptors genes upregulated in CD8 T cells (logFC > 0.5 and p-values ≤ 0.05). Dot size represents the percentage of CD8 T cells expressing each receptor gene and color indicates the average expression level.

While CellChat provides a global view of intercellular communication, it does not resolve differences arising from differential expression of L-R pairs between groups. Therefore, we applied MultiNicheNet, to identify age and diet specific communication based on differentially expressed L-R gene pairs (Browaeys et al. 2023). This analysis revealed that, during aging, most of gWAT cells uniquely signaled to CD8 T cells, primarily through MHC class Ib molecules and *Cd274* (Figure 3c and S4). In addition to MHC-1b and Cd274 mediated signaling, we also observed cell-type specific interactions: Tregs, γδT cells and endothelial cells (ECs) signaled to CD8 T cells via *Il10*, *Bmp4/6* and *Inhbb*. while ILCs, NK cells and neutrophils signaled through *Tgfb1*.

To evaluate global shifts in signaling pathways, we applied the RankNet function of CellChat, which showed age-related enrichment in IL-10, PARs (protease activated receptors), KLK (kallikrein-related peptidases), CCL (chemokines), CEACAM (adhesion molecules), and MHC-I signaling pathways (Figure 3d). Enrichment in MHC-I and IL10 signaling pathways in aged mice suggests the development of cytotoxic and exhausted phenotype of CD8 T cells (Smith et al. 2018; Raskov et al. 2021; Smith et al. 2018). Further analysis of receptor expression in aged CD8 T cells using MultiNicheNet revealed increased transcription of *Il15ra*, *Il10ra*, *Il7r*, *Ccr7*, *Acvr2a*, *Fcgr2b*, *Epha3* (Figure 3e and S5). Notably, *Il7r* and *Ccr7* are markers of naïve and central memory (CM) CD8 T cells, facilitating their homing to lymphoid structures. Given the well-established age-associated decline in naïve CD8 T cells (Goronzy et al. 2015), we hypothesize that increased expression of *Il7r* and *Ccr7* reflects the accumulation of CM CD8 T cells in aged gWAT. Overall, our cell-cell communication analysis suggests that the aged gWAT microenvironment derives CD8 T cells towards an immunosuppressive phenotype, through signaling mediated by *Cd274*, MHC-1b, *Il10*, *Bmp4/6* and *Tgfb2*.

### 2.4 CD8 T cells in aged mice exhibit a distinct phenotypic landscape

To further characterize the phenotypic diversity of CD8 T cells in gWAT, we extracted and re-clustered the CD8 expressing population from SVF (shown in Figure 2b). Dimensionality reduction and reanalysis of 2,408 CD8 T cells using UMAP revealed eight distinct clusters (Figure 4a). Cluster identities were assigned based on the expression of canonical marker genes (Figure 4c and S6). Naïve CD8 T cells were identified by high expression of stemness- and survival-associated genes, including *Lef1*, *Tcf7*, *Foxp1*, and *Il7r*, along with *Ccr7* and *Sell*. Effector memory (EM) T cells (Tem) were distinguished by high expression of memory-associated genes (*Eomes*, *Cd44*, *Ccl5*) and intermediate expression of exhaustion markers (*Pdcd1*, *Ctla4*). Two clusters (Tex1 and Tex2) represented exhausted CD8 T cells, characterized by elevated expression of *Tox*, *Nr4a2*, *Nr3c1*, *Ikzf3*, *Pdcd1*, and *Ctla4* and reduced expression of *Tcf7*. Tex1 also expressed high levels of the tissue-residency markers *Cd69* and *Cxcr6*, indicating residency phenotype acquired by these exhausted CD8 T cells. Both effector memory and exhausted CD8 T cells expressed high levels of *Gzmk*, which has earlier been associated with inflammaging (Mogilenko et al. 2021). Transcripts levels of *Gzmk* were high in both aged and obese mice (Figure S5). Clusters Tcm/vm1, Tcm/vm2, and TGzmm displayed features consistent with classical or virtual memory-like T cells. These clusters showed elevated expression of central memory (CM) markers (*Cd44*, *Ccr7*, *Sell*) and lacked *Itga4* (Cd49d), an integrin upregulated upon antigen exposure, supporting a virtual memory (VM) like identity (Hussain & Quinn 2019; Chiu et al. 2013; Clambey et al. 2008). VM CD8 T cells, which arise independently of antigen stimulation, rely on cytokines such as IL-7, IL-15, and IL-18 for maintenance, and accordingly expressed *Il2rb*, *Il4ra*, *Il7r*, and *Il18r1* (Hussain & Quinn 2019). Cluster TGzmm further expressed *Gzmm* and *Prf1*, suggesting cytotoxic function of these VM-like cells (Voskoboinik et al. 2015). Moreover, CM/VM-like CD8 T cells exhibited high expression of regulatory/inhibitory receptors including *Fcgr2b*, *Acvr2a*, *Klrc1* and *Klrd1* (Figure 4c, and S6). The Teff cluster was annotated as effector CD8 T cells based on high expression of cytotoxicity related genes *Tbx21*, *Zeb2*, *Gzmb*, *Prf1*, *Klrd1*, and *Klrg1*. As expected, HFD-fed mice and ND-fed aged mice displayed elevated frequencies of exhausted CD8 T cells (Tex1). However, age-related changes were more pronounced in Tcm/vm1, Tcm/vm2, and TGzmm clusters compared to either ND- or HFD-fed young mice (Figure 4b).

**Figure 4.**
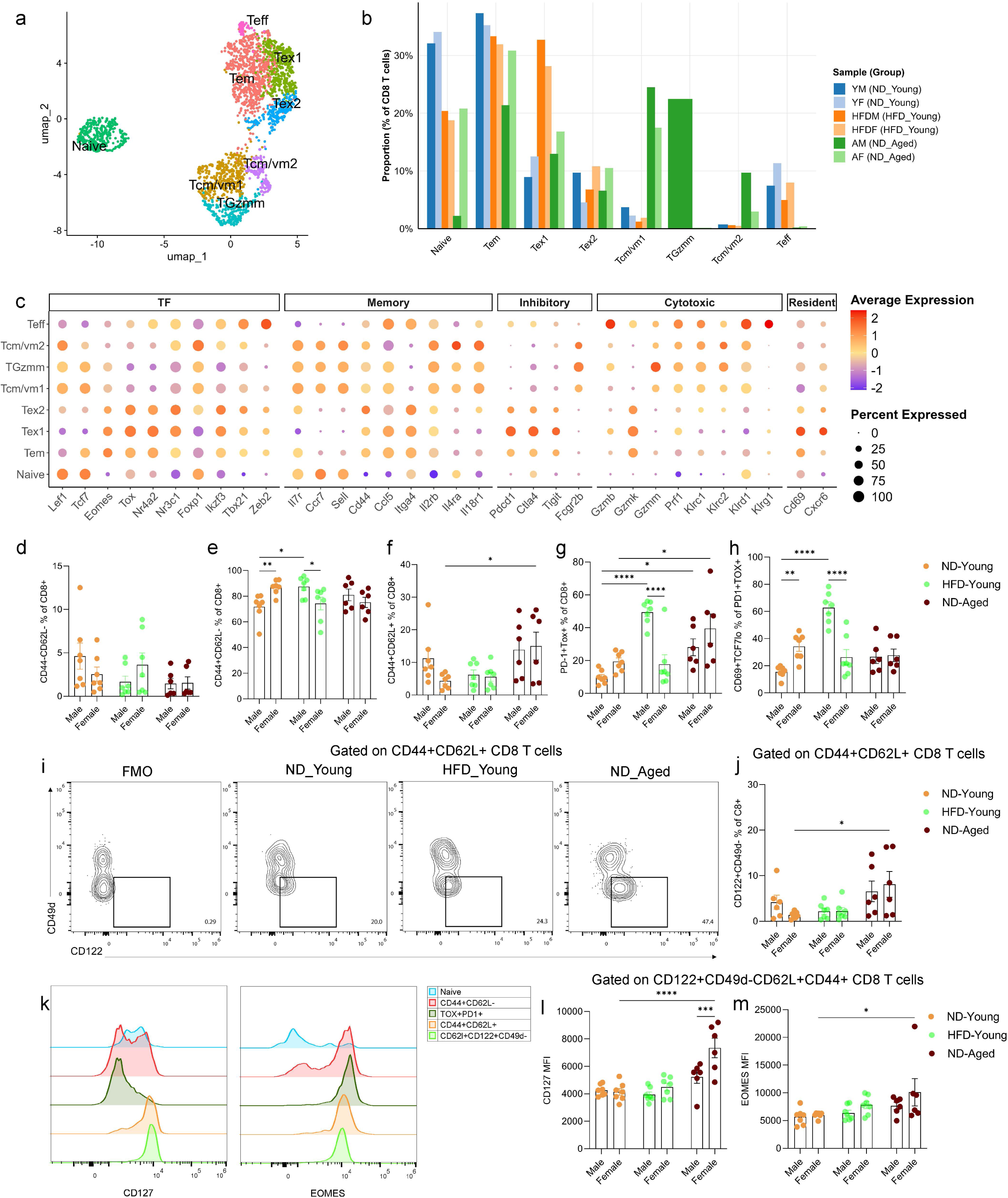
scRNA-seq analysis of CD8 T cells. (a) UMAP plot showing 2,408 CD8 T cells following subsetting from SVF of ND-fed young, HFD-fed young, and ND-fed aged mice, colored by annotated cell-types. (b) Bar plot representing the proportions of CD8 T cell subsets relative to total CD8 T cells. (c) Dot plot displaying expression of marker genes used to define CD8 T cell subsets. Dot size represents the fraction of cells expressing each marker gene and color indicates the average expression across subsets. Bar plots showing frequencies of (d) CD44-CD62L-, (e) CD44+CD62L-, (f) CD44+CD62L+, and (g) PD-1+Tox+ of CD8 T cells measured using flow cytometry across different groups. (h) Bar plot showing frequencies CD69+TCF7lo cells within PD-1+Tox+ CD8 T cells. (i) Contour plot representing gating strategy (j) Bar plot showing frequency of CD122+CD49d− virtual memory CD8 T cell across different groups. (k) Histograms illustrating the expression of CD127(Il7r) and Eomes in various CD8 T cell subsets. Bar graphs representing median fluorescent intensity of (l) CD127, and (m) Eomes on CD122+CD49d-CD62L+CD44+ CD8 T cells across different groups. Data are presented as mean ± SEM. Statistical significance was determined using two-way ANOVA followed by Tukey’s HSD post hoc test. Only significant comparisons are shown between sex of each group and with respect to ND-fed young mice. N = 6-7 mice per group. P-values < 0.05 were considered significant. p< 0.05 (*), p<0.01 (**), p<0.001 (***), p<0.0001 (****). Gating strategy used to plot the cell types is shown in Figure S8.

To validate the scRNA-seq findings, we performed multicolor flow cytometry on SVF isolated from gWAT. Although naïve (CD44-CD62L-) CD8+ T cell infiltration was reduced in gWAT of ND-fed aged mice compared to ND-fed young mice, this difference was statistically non-significant (Figure 4d). As expected, HFD-fed male mice exhibited significant accumulation of PD-1+ TOX+ and PD-1+ TOX+ CD69+ TCF-7lo CD8+ T cells, aligning with the exhausted phenotype seen in scRNA-seq data (Figure 4g, and 4h). HFD-fed males also showed a significant increase in EM (CD44+CD62L-) CD8+ T cells. Since exhausted CD8+ T cells also express a CD44+CD62L− phenotype (Figure 4c), direct gating of PD-1+ cells on CD8+ T cells in flow cytometry may lead to overestimation of EM population. In aged mice, we observed a significant accumulation of PD-1+ TOX+ CD8+ T cells. However, the frequency of PD-1+ TOX+ CD69+ TCF-7lo CD8+ T cells was elevated in aged males, without reaching significance, suggesting that HFD exerts a more robust effect on the transition to exhaustion than physiological aging. Notably, aging was associated with significant increases in CM (CD44+CD62L+) and VM-like (CD44+CD62L+CD122+CD49d-) CD8+ T cell subsets (Figure 4f, 4i, and 4j), consistent with scRNA-seq results (Figure 4b). These VM-like CD8+ T cells expressed higher levels of CD127 (*Il7r*) and maintained Eomes expression (Figure 4k). Compared with young controls, aged VM-like CD8+ T cells exhibited elevated levels of both CD127 and Eomes compared with young controls (Figure 4l, and 4m). Although these protein expression changes were more pronounced in aged female mice, expansion of VM-like CD8+ T cells during aging suggests a unique and potentially critical role for these cells in adipose tissue during aging.

### 2.5 Trajectory analysis reveals dysregulated differentiation stages of aged CD8 T cells

To explore the differentiation dynamics of CD8 T cells in aged gWAT, we performed RNA velocity analysis, Monocle trajectory inference, and pseudotime analysis. RNA velocity estimates future cell states based on spliced and unspliced mRNA ratios, while Monocle infers differentiation trajectories based on gene expression changes along pseudotime (Trapnell et al. 2014; Bergen et al. 2020; La Manno et al. 2018). Both methods revealed similar trends in the developmental trajectories (Figure 5a, 5b, and 5c). Monocle inferred pseudotime trajectories suggested a progression from the Tcm/vm1 cluster towards the Tem cluster, consistent with prior reports of homeostatic, antigen independent proliferation from CM to EM cells (Bouneaud et al. 2005; Geginat et al. 2003; Geginat et al. 2001). Both RNA velocity and Monocle predicted that memory subsets could subsequently transition into a Tcm/vm2 phenotype. Tem cells could differentiate into exhausted subsets (Tex1 and Tex2), which could further transition into a Tcm/vm2 phenotype. Additionally, cluster TGzmm appeared to be differentiated from Tcm/vm1, both TGzmm and Tcm/vm1 cells at later stages could converge toward the Tcm/vm2 phenotype.

**Figure 5.**
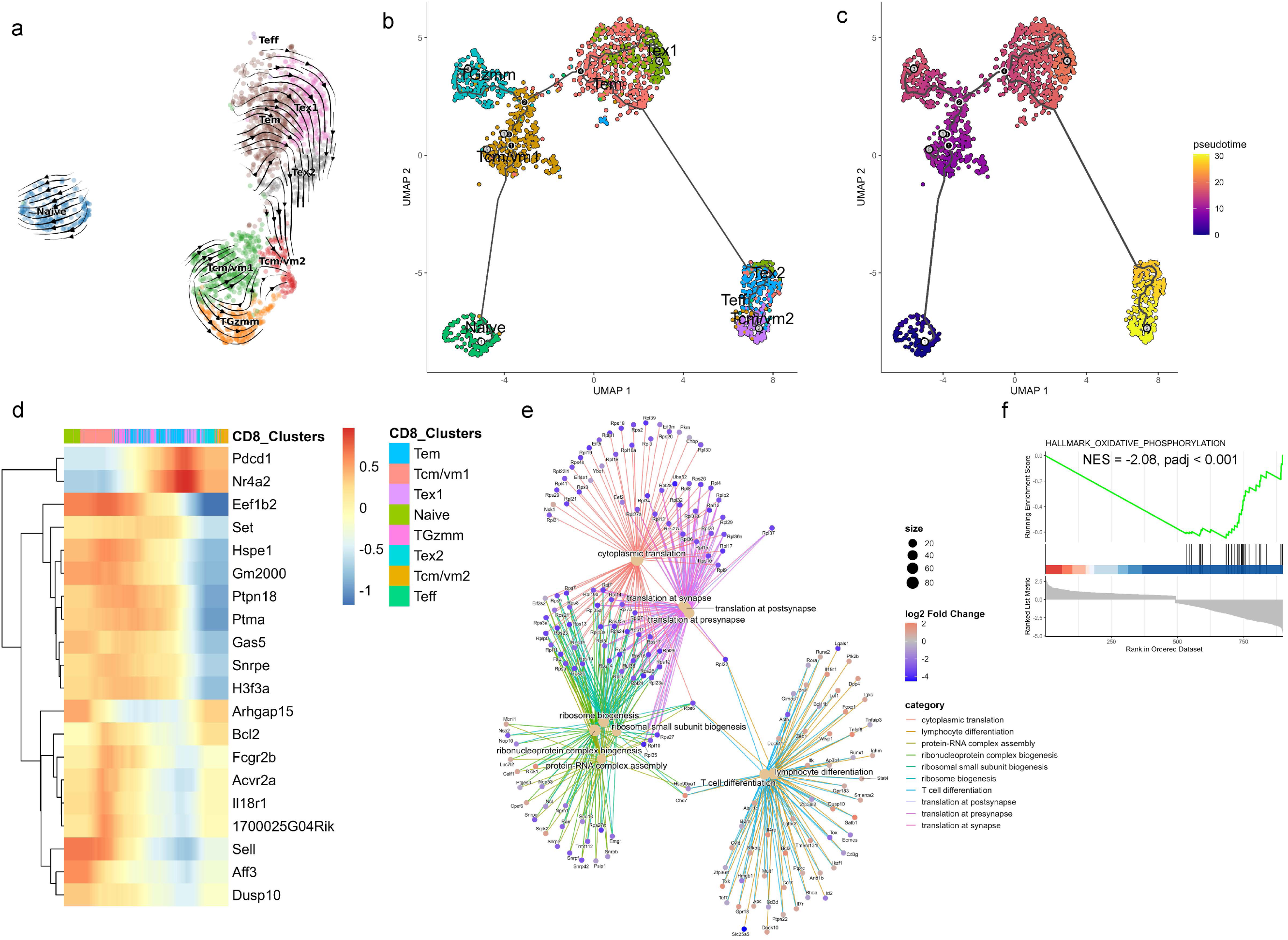
RNA velocity and trajectory analysis of CD8 T cells. (a) UMAP plot showing trajectories inferred from RNA velocity, (b) Trajectory constructed using Monocle, and (c) Pseudotime projection of CD8 T cells from ND-fed aged mice. (d) Heatmap showing the dynamic changes in gene expression along pseudotime, (e) cnetplot illustrating functional relationships among the top 10 significant Gene Ontology (GO) terms and their associated genes in the Tcm/vm2 cluster. Node size reflects p-value of each GO term, and color indicates the log2 fold change in gene expression. (f) Gene set enrichment analysis (GSEA) enrichment plot for hallmark oxidative phosphorylation gene set in the Tcm/vm2 cluster.

To further define these differentiation pathways, we plotted the top 20 differentially expressed genes along the pseudotime in a heatmap (Figure 5d). Cells at the earliest pseudotime points exhibited high expression of *Sell* and *Dusp10*, indicative of a naïve phenotype. The transition towards CM/VM-like phenotype was marked by increased expression of *Fcgr2b*, *Acvr2a*, and *Il18r1*, alongside sustained *Sell* expression. Differentiation of TGzmm cells from Tcm/vm1 was characterized by high expression of *Fcgr2b* and intermediate levels of *Sell* and *Il18r1* (Figures 4c, 5d). Acquisition of exhaustion phenotype (Tex1) was associated with upregulation of *Pdcd1* and *Nr4a2*. Differentiation into the Tcm/vm2 subset was marked by downregulation of genes involved in protein synthesis (*Eef1b2*, *Hspe1*, *Snrpe*, *H3f3a*), intermediate expression of *Fcgr2b*, *Dusp10*, and *Il18r1*, and high expression of *Bcl2*. This transcriptional profile suggests that Tcm/vm2 cells possess reduced translational activity while maintaining survival potential and immunological responsiveness. To functionally characterize the Tcm/vm2 population, we performed overrepresentation analysis and Gene Set Enrichment Analysis (GSEA) which revealed significant defects in protein translation, ribosome biogenesis, and oxidative phosphorylation pathways (Figure 5e, and 5f). At the same time, expressions of *Lef1*, *Foxp1*, and *Dusp10* suggest a resting or quiescent phenotype, whereas expressions of cytokine receptors (*Il4r*, *Il7r*, *Il18r1*) implied that these cells could respond to cytokines (Figure 4c, 5d, and 5e). Overall, trajectory analysis not only mapped the differentiation pathways of CD8 T cells in aged adipose tissue but also identified key regulatory genes like *Fcgr2b*, *Acvr2*, *Il18r1* whose expression dynamics may govern fate decisions within the CM/VM-like CD8 T cells during aging.

### 2.6 Accumulation of Fcgr2b expressing CM/VM-like CD8 T cells in aged gWAT drives inflammation

The inhibitory receptor *Fcgr2b* has been previously implicated in regulating CD8 T cells, with its loss associated with an accumulation of effector cells (Morris, Farley, et al. 2020). In our dataset, pseudotime trajectory analysis suggested a regulatory role for *Fcgr2b* in CD8 T cell differentiation, with expression predominantly confined to CV/VM-like (*Itga4*-) CD8 T cells rather than EM cells (Figure 4b, 6a, and 6b). Consistent with scRNA-seq data, ND-fed aged mice showed a significant increase in the frequencies of Fcgr2b/CD32b+CD49d− cells within the CD44+CD62L+ CD8+ T cells compared to both ND- and HFD-fed young mice (Figure 6c, 6d and S5). Moreover, CM/VM-like CD8+ T cells demonstrated elevated expression of GZMM relative to other CD8 T cell subsets (Figure 4c, 6e, and 6f). Notably, aging was associated with a significant upregulation of GZMM, with ND-fed aged males exhibited higher levels of GZMM compared to ND-fed aged females, indicating sex-specific differences in GZMM production (Figure 6g, and S5). Ligand-receptor analysis revealed that aged CD8 T cells could interact with multiple cell types, including macrophages and fibroblasts, via distinct signaling axes (Figure 6h, and S7). Given association between GZMM and inflammation (Shan et al. 2020; Baschuk et al. 2014; Anthony et al. 2010), we next explored its functional role *in-vitro* on mouse macrophages and fibroblasts. Mouse bone marrow-derived macrophages (BMDMs), both unprimed and LPS-primed (Figure 6i, and 6j), and cycling and senescent mouse embryonic fibroblasts (MEFs) (Figure 6i, and 6k), were stimulated with recombinant mouse GZMM. In all cell types, GZMM treatment significantly increased the secretion of pro-inflammatory proteins, including IL-6, CXCL1, and CCL2. These findings suggest that aged CD44+CD62L+Fcgr2b+CD49d-CD8+ T cells, while retaining cytotoxic potential, may also contribute to adipose tissue inflammation via GZMM secretion. This expands the established paradigm of granzyme-mediated inflammation, previously attributed to GZMK-expressing exhausted CD8 T cells, by suggesting that both GZMK and GZMM may mediate inflammation in the adipose tissue of aged mice through distinct CD8 T cell subsets (Mogilenko et al. 2021).

**Figure 6.**
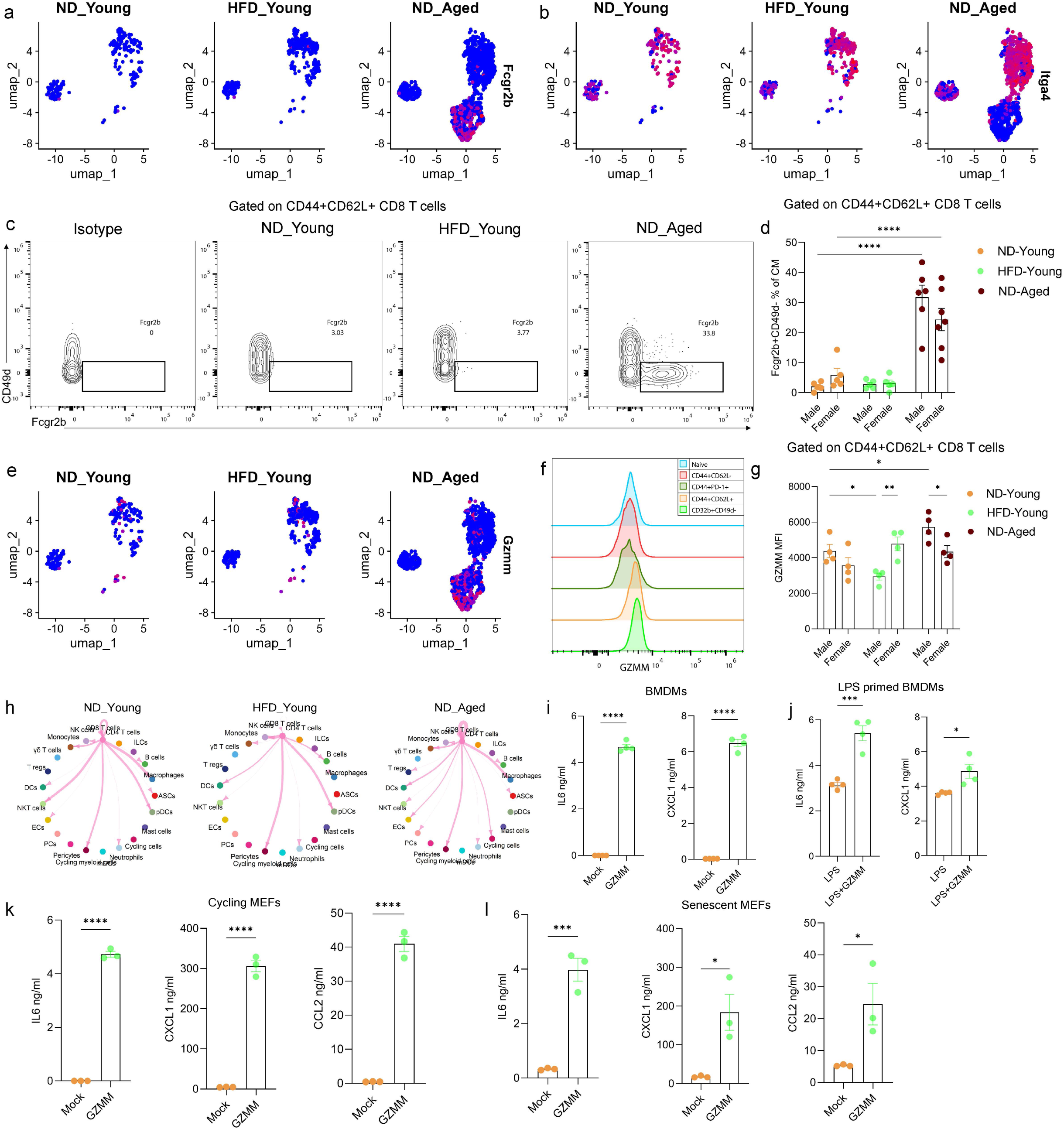
Accumulation of Fcgr2b expressing cells in the gWAT of aged mice. UMAP plot showing expression of (a) *Fcgr2b* and (b) *Itga4* in CD8 T cell from ND-fed young, HFD-fed young, ND-fed aged mice. (c) Representative flow cytometry plots displaying the gating strategy for identifying Fcgr2b+CD49d− cells within CD44+CD62L+ CD8 T cells. (d) Bar graph showing frequency of Fcgr2b+CD49d− within CD44+CD62L+ CD8 T cells (N=5-7). (e) UMAP plot showing expression of *Gzmm* in CD8 T cells among different groups. (f) Histogram showing GZMM expression in CD8 T cell subsets measured using flow cytometry. (g) Bar plot representing median fluorescent intensity (MFI) of GZMM in CD44+CD62L+ CD8 T cells across different groups (N=4). Statistical significance was determined using two-way ANOVA followed by Tukey’s HSD post hoc test. Only significant comparisons are shown between sex of each group and with respect to ND-fed young mice. (h) Circle plot depicting outgoing signals from CD8 T cells to other cell types in SVF based on L-R expression,ss analyzed using Cellchat. Each node represents a cell type, and edge width reflects interaction strength. Quantification of IL6, and CXCL1 levels in supernatants of (i) unprimed and (j) LPS primed bone marrow derived macrophages (BMDMs) following overnight stimulation with GZMM, measured by ELISA. Quantification of IL6, CXCL1, and CCL2 in the supernatants of (k) cycling and (l) senescent mouse embryonic fibroblast (MEFs) following stimulation with GZMM for 48 hours and 24 hours, respectively, measured by ELISA. Statistical significance was determined using unpaired t-test. Data are presented as mean ± SEM. P-values < 0.05 were considered significant. p< 0.05 (*), p<0.01 (**), p<0.001 (***), p<0.0001 (****).

## 3 Discussion

Distinguishing the specific cellular changes associated with aging and obesity is essential for understanding their individual contributions to disease and for developing targeted interventions. Using scRNA-seq approach, we delineated the heterogeneity of VAT during physiological aging and HFD-induced obesity. We observed that aging was associated not only with a marked accumulation of CD8 T cells in gWAT, but also with increased interaction strength between CD8 T cells and SVF cells, primarily mediated MHC-I and IL10 signaling. While both aging and obesity led to expansion of exhausted CD8 T cells, aging uniquely promoted accumulation of a phenotypically distinct population of CD8 T cells resembling VM-CD8 T cells. These cells expressed canonical memory-associated markers *CD44*, *Sell*, *Il7r*, *Il2rb,* and exhibited reduced expression of *Itga4*, a VM phenotype. Notably, this subset also expressed *Fcgr2b*, *Acvr2a* along with *Gzmm*, suggesting a potential role in immune regulation, cytotoxicity and inflammation. These findings suggest that aging uniquely shapes the adipose tissue microenvironment to promote a VM-like CD8 T cell phenotype with both regulatory and cytotoxic potential.

Both aged and obese mice exhibited increased signaling toward CD8 T cells. The majority of this signaling was mediated by engagement of MHC-Ib molecules in both conditions. Although MHC-I typically present self-peptides, the development of a senescent phenotype in the adipose tissue cells during aging and obesity could explain the increased MHC-I signaling compared to ND-fed young mice (Zhang et al. 2024; Marin et al. 2023; Ou et al. 2022; Pereira et al. 2019; Palmer et al. 2019). MHC-Ib molecules can interact with either inhibitory or activating receptors on CD8 T cells, and the outcome of this interaction likely shapes their functional response. In our dataset, CM/VM-like CD8 T cells expressed high transcript levels of both inhibitory receptor Klrc1(NKG2A) and activating receptor Klrc2 (NKG2C). Given that NKG2A exhibits higher binding affinity to MHC-Ib molecules than NKG2C, these interactions may favor inhibitory signaling and functional suppression of CD8 T cells (Wu et al. 2023; Pereira et al. 2019; Béziat et al. 2011). Notably, most cell types, such as B cells, DCs, ILCs, Tregs, γδT cells, PCs, etc., in the gWAT of aged and obese mice exhibited increased MHC-I signaling compared to young controls, with the exception of Pdgfra-expressing ASCs. This suggests that ASCs may be vulnerable to CD8 T cells-mediated cytotoxicity, although this requires further validation.

In addition to MHC-I mediated signaling, CD8 T cells in gWAT also received signals via *Cd274* (PD-L1), *Il10*, *Bmp4/6*, and *Inhbb* ligands. Correspondingly, elevated expression of their receptors, *Epha3*, *Il10ra*, *Acvr2a*, were observed on CD8 T cells of aged mice. Both PD-L1 and IL10 are known to induce an immunosuppressive state in CD8 T cells (Cha et al. 2019; Smith et al. 2018). In our analysis, *Epha3* and *Il10ra* were selectively expressed in effector and exhausted CD8 T cell subsets, supporting their role in promoting T cell dysfunction. Although the role of *Epha3* in CD8 T cells is not well defined, it has been reported to be upregulated in malignant T cells (Maddigan et al. 2011; Smith et al. 2004).

Notably, we found *Acvr2a* was highly expressed in the *Klrc1*-expressing CM/VM-like CD8 T cell cluster. As *Acvr2a* can signal through both SMAD1/5 and SMAD2/3 pathways, the relative balance of *Bmp4*, *Inhbb*, and *Tgfb1* signaling in the aged adipose tissue microenvironment may influence CD8 T cell fate by either promoting a dysfunctional state or inducing stemness (Hu et al. 2025; Saadey et al. 2023; Pinjusic et al. 2022; Appiah Adu-Gyamfi et al. 2020; Olsen et al. 2015). Increased frequencies of VM CD8 T cells during aging have been well documented in the peripheral blood, spleen and lymph nodes (Davenport et al. 2019; Quinn et al. 2018; Chiu et al. 2013; Hussain et al. 2023; Borsa et al. 2021; Quinn et al. 2020; Hussain & Quinn 2019). In our dataset, we have further demonstrated their accumulation in gWAT of aged mice. Both CM and VM share multiple features including expression of *Sell* (CD62L), *Cd44*, *Il7r*, *Eomes*, but can be distinguished by CD49d, which is absent on VM CD8 T cells (Hussain & Quinn 2019). Accordingly, we identified VM CD8 T cells as CD122+CD49d− within the CD44+CD62L+ CD8 T cell population. The accumulation of CD49d− VM CD8 T cells in aged adipose tissue may be driven by cytokine mediated homeostatic expansion (Quinn et al. 2018; Chiu et al. 2013; Renkema et al. 2014).

Studies from the past decade have demonstrated the *Fcgr2b* expression on memory CD8 T cells, contrary to the thought of their expression only on B cells and on innate immune cells (Morris, Pinelli, et al. 2020; Morris, Farley, et al. 2020; Starbeck-Miller et al. 2014; Nimmerjahn & Ravetch 2008). In the present study, we demonstrated higher expression levels of *Acvr2a* and *Fcgr2b* on CM/VM-like CD8 T cells of aged mice, which showed a strong association with pseudotime trajectories, suggesting their involvement in CD8 T cell fate decision. Fcgr2b is the only inhibitory Fc receptor that signals through an immunoreceptor tyrosine based inhibitory motif (ITIM) in its cytoplasmic domain (Getahun & Cambier 2015). Morris et al. reported that *Fcgr2b* is predominantly expressed by effector memory (CD44+CD62Llo) CD8 T cells, and its engagement by Fgl2 can induce apoptosis and limit their expansion (Morris, Farley, et al. 2020). Additional studies suggest that *Fcgr2b* expression on effector CD8 T cells impair responsiveness to anti-PD-1 therapy, whereas its deletion enhances CD8 T cell stemness (Ku et al. 2024; Bennion et al. 2023). Microbial stimulation has been shown to induce Fcgr2b expression on memory CD8 T cells (Morris, Pinelli, et al. 2020; Starbeck-Miller et al. 2014), its engagement by preexisting antigen-antibody complexes has been reported to limit expansion of memory cells during homologous rechallenge (Starbeck-Miller et al. 2014). More recently *Fcgr2b* has been detected on aged regulatory CD8 T cells, suggesting broader immunoregulatory functions during aging (Srinivasan et al. 2025).

In our scRNA-seq data, *Fcgr2b* transcripts were enriched in CM/VM-like CD8 cells compared to EM CD8 T cells. Expression of *Fcgr2b,* as well as proportion of *Fcgr2b* expressing cells, was highest in the TGzmm cluster. Flow cytometry further confirmed age-related accumulation of Fcgr2b+CD49d− negative CD8 T cells in gWAT. Both previous reports and our trajectory analysis support the regulatory role of Fcgr2b in CD8 T cell fate during aging. Moreover, age-associated accumulation of IgG, a *Fcgr2b* ligand, has been reported in visceral adipose tissue (Yu et al. 2024). While engagement of IgG-Fcgr2b could theoretically promote apoptosis, these cells expressed high levels of Bcl2, suggesting a mechanism for survival under pro-apoptotic pressure. Interestingly, these cells also expressed Fcgrt, the neonatal Fc receptor, further supporting a role of IgG in modulating CD8 T cell function in gWAT of aged mice. RNA velocity indicated that *Fcgr2b* expressing TGzmm cells may differentiate into dysfunctional state (Tcm/cm2) suggest the engagement of Fcgr2b in aged mice. Consistent with the previous report, we also observed that *Fcgr2b* expressing cluster exhibited a cytotoxic profile (Bennion et al. 2023). These data suggest that the *Fcgr2b* expressing TGzmm cluster represents a transitional cell state capable of inducing cytotoxicity and granzyme M production.

The accumulation of CD8 T cells has been linked to adipose tissue inflammation (Nishimura et al. 2009). More recently, age-associated *Gzmk* expressing CD8 T cells (Taa) has been shown to induce inflammatory response in fibroblast cells (Mogilenko et al. 2021). In our dataset, *Gzmk* expression was elevated in both aged and obese mice CD8 T cells, while *Gzmm* expression was selectively increased in aged mice CD8 T cells. Although granzymes are well known to mediate cytotoxicity, emerging evidence suggests that they can also regulate inflammation (Aubert et al. 2024; Zheng et al. 2023; Shan et al. 2020; Baschuk et al. 2014; Anthony et al. 2010). We found that GZMM treatment was sufficient to induce proinflammatory cytokine release from both mouse fibroblasts and macrophages. Although *Gzmk* expressing cells make up a large fraction of CD8 T cells (∼50% proportion of CD8 T cells, including Tem, Tex1 and Tex2 clusters) which may induce inflammation in aged adipose tissue, *Gzmm* expressing cells (∼20% TGzmm cluster) could further contribute exacerbate the inflammatory burden.

Collectively, we have shown that VM-like CD8 T cells accumulate in visceral adipose tissue during aging. Their differentiation trajectory appears to involve signaling through *Acvr2a* and *Fcgr2b*. During transition towards dysfunctional states, these cells acquire cytotoxic potential, marked by *Gzmm* expression, which may contribute to cytotoxicity and adipose tissue inflammation. Future studies will be required to determine the specific role of these cells in regulating cytotoxicity and inflammation in aged adipose tissue.

This study has some limitations. First, we were unable to validate Acvr2a expression using flow cytometry because of the lack of specific monoclonal antibody suitable for flow cytometry. However, we observed high levels of Acvr2a transcripts in CM/VM clusters, suggesting potential significance of this receptor in CD8 T cells during aging. Second, we did not utilize Fcgr2b knockout mouse to directly assess the functional significance of Fcgr2b expressing cells in aging or their role in modulating CD8 T cell fate. However, our findings highlight the presence of Fcgr2b expressing VM CD8 T cells in aged mice, and their potential functional significance. Future studies are needed to elucidate the function of these cells in the context of aging and age-related diseases.

## 4 Conclusion

Our study provides a comprehensive single-cell analysis of SVF isolated from gWAT of aged and obese mice of both sexes. We highlighted key differences in the cellular heterogeneity of gWAT between aging and obesity. Using cell-cell interaction and trajectory analysis, we identified *Acvr2a* and *Fcgr2b* may modulate CD8 T cell differentiation in aged adipose tissue. We further show that *Fcgr2b* expressing cells exhibit a cytotoxic profile and may contribute to Gzmm-dependent inflammatory signaling. These findings provide new insights into VM-like CD8 T cells in aged adipose tissue and suggest *Acvr2a* and *Fcgr2b* as potential modulators of their differentiation and functional state during aging.

## 5 Methods

### 5.1 Mice and Diet

C57BL/6J male and female mice (young: 1-2 month; aged: 18-21 month) were obtained from Jackson Laboratory. Mice were housed in a specific pathogen-free (SPF) facility at the University of Michigan. Young mice were fed a high-fat diet (HFD) (42% Kcal from fat, Inotiv; TD.88137) for an additional 12 weeks, while the rest of the young and aged mice were maintained on a normal diet (ND) for the same time period. All the experiments were approved by the Unit of Laboratory Animal Medicine, University of Michigan, under animal protocols PRO00010459 and PRO00012394 and were performed accordingly.

### 5.2 Glucose and Insulin Tolerance Tests (GTT and ITT)

For glucose tolerance tests (GTT), mice were fasted overnight and intraperitoneally (i.p.) injected with glucose at a dose of 2.0 g/kg body weight. For insulin tolerance tests (ITT), mice were fasted for 6 hours prior to i.p. injection of Humulin R (Eli Lilly & Co) at 0.8 U/kg body weight. Blood glucose levels were measured using Clarity BG1000 blood glucose monitoring system (Clarity Diagnostics) at baseline and at intervals of 15-30 minutes for up to 2 hours post-injection.

### 5.3 Stromal Vascular Fraction (SVF) isolation

Mice were euthanized using CO_2_ inhalation, and gonadal white adipose tissue (gWAT) was dissected. Tissue was finely minced and digested in 0.8 mg/ml Collagenase II (Worthington Biochemical) buffer containing 3% BSA, 1X penicillin/streptomycin, 1.2 mM CaCl₂, 1 mM MgCl₂, 0.8 mM ZnCl₂, and 15 mM HEPES for 40 minutes at 37°C with constant agitation. The digested suspension was centrifuged at 600g for 10 minutes at 4°C, and the SVF was filtered through 70 µm and 40 µm strainers. Red blood cell (RBC) lysis was performed as per manufacturer’s instructions (eBioscience), followed by washing and resuspension in RPMI1640 supplemented with 10% FBS and 1X penicillin/streptomycin. Fresh cells were processed for single cell RNA sequencing and remaining cells were cryopreserved.

### 5.4 Single-cell RNA sequencing (scRNA-seq)

SVF cells were stained with Zombie Aqua live/dead dye (Biolegend) for 10 minutes at room temperature. Cells were sorted using a Bigfoot Spectral Cell Sorter (Thermo Fisher Scientific). Sorted cells were resuspended in RPMI medium with 10% FBS, equal number of cells per group were pooled and submitted to the University of Michigan Advanced Genomics Core for 3′ library preparation and sequencing. Briefly, cell counts were obtained using the Luna-FX7 Cell Counter (LogosBio). Libraries were prepared using the 10X Genomics Chromium Controller with 3′ v3.1 chemistry and Feature Barcoding technology for Cell Multiplexing, according to the manufacturer’s instructions (10X Genomics). Library quality was assessed on the LabChip GXII HT (PerkinElmer), and concentrations were determined using Qubit (Thermo Fisher). Pooled libraries were sequenced using paired-end 28×10×10×151 bp reads on the Illumina NovaSeq XPlus. Demultiplexed FASTQ files were generated using Bcl2fastq2 (Illumina), and count matrices were generated using the CellRanger pipeline (10X Genomics) (Zheng et al. 2017).

### 5.5 scRNA-seq analysis

Data were analyzed using Seurat v5.1.0 (Butler et al. 2018). Cells were filtered to retain those with ≥500 UMIs, ≥250 genes, log10(genes per UMI) > 0.8, and mitochondrial gene content <15%. Genes expressed in fewer than 10 cells were excluded. Doublets were detected using scDblFinder v1.16.0 (Germain et al. 2022) and further removed. Each dataset was normalized, and 4,000 variable features were identified using the “vst” method. Integration was performed using SelectIntegrationFeatures, FindIntegrationAnchors, and IntegrateData. Variables including gene count, mitochondrial ratio, and cell cycle scores (S and G2M) were regressed during scaling. PCA was performed using 60 principal components (PCs). UMAP was applied on the top 40 PCs, guided by elbow plot analysis. Clustering was performed at a resolution of 1.2. Cell clusters were annotated using SingleR v2.4.1, referencing the ImmGen and MouseRNAseq datasets from the cellDex v1.12.0 package (Aran et al. 2019). Annotations were finally refined manually based on top marker genes identified via FindAllMarkers.

### 5.6 Cell-Cell communication analysis

Intercellular communication in the adipose SVF was analyzed using CellChat v2.1.2 and MultiNicheNet v2.0.1 (Jin et al. 2025; Browaeys et al. 2023). Clusters with less than 10 cells in any group were removed from the analysis. Ligand-receptor (L-R) interactions were analyzed across various cell types by calculating communication probabilities between different cells using computeCommunProb(), and filtering out communication involving less than 10 cells using filterCommunication().

For focused analysis of signaling input to CD8 T cells, MultiNicheNet was employed to identify L-R pairs based on differentially expressed genes (DEGs) from multi-group data. Cell types with <10 cells per group were excluded and DEGs with log₂ fold change ≥ 0.5 and adjusted p-value ≤ 0.05 were used to define the top 100 L-R interactions across conditions.

### 5.7 CD8 T cell subset analysis

CD8a expressing cells were subsetted into a new Seurat object and subjected to standard preprocessing and clustering as described earlier. UMAP dimensionality reduction was performed using 15 PCs and clusters were resolved at a resolution of 0.6. CD8 T cell clusters were manually annotated based on the expression of canonical makers. Naïve = (*Ccr7*+, *Lef1*+ *IL7r*+ *Sell*+ *Cd44*-*Ccl5*-), Tcm/vm-like cells (*Ccr7*+, *Lef1*+ *IL7r*+ *Sell*+ *Cd44*+ *Ccl5*+ *Il2rb*+ *Il18r1*+ *Itga4*-), Tem (*Cd44*+ *Sell*-*Tcf7*+ *IL7r*+ *Ccl5*+), Tex (*Pdcd1*, *Tox*, *Nr4a2*, *Nr3c1*), Teff (*Tbx21*+, *Zeb2*+ *Gzmb*+ *Prf1*+ *Klrg1*+). Markers expression was visualized using Dotplot function and clusters proportions were calculated relative to the total CD8 T cells. Enrichment analysis was performed on differentially expressed genes from the Tcm/vm2 cluster using the clusterProfiler (4.10.1) (Yu et al. 2012). Gene Ontology (GO) was performed using function enrichGO() and top 10 significant GO terms were visualized using cnetplot(). Gene set enrichment analysis (GSEA) was performed using Hallmark pathways from MSigDB using msigdbr (10.0.1).

### 5.8 RNA velocity and Monocle pseudotime analysis

RNA velocity analysis was performed using Velocyto (v0.17.17) and scVelo (v0.3.3) (La Manno et al. 2018; Bergen et al. 2020). Individual loom files were generated from the CellRanger output using the velocyto and subsequently merged using loompy.combine(). The combined loom file was loaded into Scanpy (v1.11.0) (Wolf et al. 2018), and CD8 T cells were subset for further analysis using metadata, cell barcodes and UMAP coordinates exported from Seurat. RNA velocity was computed using scv.tl.velocity(mode=’stochastic’) and visualized on CD8 UMAP embeddings using scv.pl.velocity_embedding_stream, with cells colored by CD8 clusters.

Trajectory inference was performed using Monocle3 (1.3.7) (Trapnell et al. 2014). Seurat objects were converted into cell_data_set format and size factors were estimated with estimate_size_factors(). Cells were clustered and trajectories were learned using learn_graph(). Root cells were defined on naïve CD8 T cell and pseudotime was assigned via order_cells(). Cells were visualized by pseudotime using plot_cells(). To identify genes dynamically expressed along pseudotime, we used the graph_test() function with the principal_graph. The top 20 pseudotime-associated genes were visualized in a heatmap using pheatmap (1.0.12), with cells ordered by pseudotime and annotated by CD8 cluster identity.

### 5.9 Flow cytometry

Cryopreserved SVF cells were thawed and 0.5-2 million cells per sample were used for staining. Cells were washed and stained for live/dead staining using Zombie aqua (Biolegend). Fc receptor blocking was performed using TruStain FcX^TM^ (anti-mouse CD16/32, Biolegend) antibody according to manufacturer instruction, except for samples stained for Fcgr2b/CD32b, where Fc blocking was omitted. Surface staining was carried out at 4°C for 30 minutes. Following surface staining, cells were washed, fixed and permeabilized using eBioscience™ Foxp3 staining buffer (Thermo) and stained for intracellular proteins. Data acquisition was performed on ID7000 (Sony) and unmixed FCS files were analyzed using FlowJo v10.10.0 (FlowJoLLC). Anti-mouse BV421 CD69 (H1.2F3), BV605 CD44 (IM7), BV711 CD62L (MEL-14), BV785 NK1.1 (PK136), BV785 CD19 (6D5), BV785 F4/80 (BM8), BV785 CD14 (Sa14-2), BV785 CD11c (N418), AF488 CD127 (A7R34), AF700 CD4 (RM4-5), APC-Fire750 PD-1 (29F.1A12), APC-Fire810 CD8 (53-6.7) were purchased from Biolegend; BUV395 CD3 (17A2), BUV496 CD45 (30-F11), BUV737 CD49d (9C10(MFR4.B)), BB700 CD122 (TM-β1) were purchased from BD Biosciences; PE Tox (TXRX10), PE Fcgr2b/CD32b (AT130-2), PE-eFluor610 Eomes (Dan11mag) were purchased from Thermo; AF647 TCF1/TCF7 (C63D9) from Cell Signaling and GZMM (MBS2042310) from MyBioSource.

### 5.10 Granzyme M in-vitro stimulation

Bone marrow derived macrophages (BMDMs) were thawed and rested overnight in DMEM supplemented with 10% FCS, 20% L929-conditioned media. BMDMs were then primed with lipopolysaccharide (LPS, 10ng/ml) for 3 hrs at 37°C. Both primed and unprimed BMDMs were washed and subsequently stimulated with mouse recombinant granzyme M (GZMM, MyBioSource; 100ng/ml) 2% FCS, 20% L929-conditioned media for 24 hrs at 37°C.

Senescence was induced in mouse embryonic fibroblasts (MEFs) as described previously (Mogilenko et al. 2021). Briefly, MEFs were cultured in 10% FCS supplemented DMEM up to 70% confluency, followed by treatment with 0.1µM doxorubicin for 24 hrs. The media was then replaced with fresh DMEM, and cells were cultured for an additional 24 hrs before being treated again with 0.1µM doxorubicin (Sigma) for another 24 hrs. MEFs were then incubated in fresh media for 7 days for senescence induction. Cycling and senescent MEFs were subsequently treated with GZMM (100ng/ml) with DMEM and without FCS for 48 hrs and 24 hrs, respectively. Supernatants collected from BMDMs and MEFs were used for quantitative estimation of IL6, CXCL1 and CCL2 by DuoSet ELISA kits, following the manufacturer’s instructions (R&D Systems).

### 5.11 Statistical analysis

Data are presented as mean ± standard error of mean (SEM). Statistical significance between groups and sexes was accessed using two-way ANOVA followed by Tukey’s HSD post hoc test. An unpaired t-test was used to compare GZMM-treated and untreated samples. P-values < 0.05 were considered significant. Significance levels are indicated in the figure legends as follows p<0.05 (*), p<0.01 (**), p<0.001 (***), p<0.0001 (****).

## Authors contributions

Study design: A.K and R.Y

Experiments: A.K and M.O’B

Data acquisition and analysis: A.K

Writing: A.K

Reviewing and editing: A.K, and R.Y

Funding: R.Y, and V.B.Y

## Acknowledgements

We thank Vern B. Carruthers (Microbiology & Immunology, University of Michigan) for providing MEFs, Yifan Wang (Microbiology & Immunology, University of Michigan) for providing BMDMs as well as reagents and suggestions for macrophage stimulation experiment, and Advanced Genomics Core (University of Michigan) for performing Library prep and next-generation sequencing. BioRender was used to create schematic and ChatGPT for English correction and text clarity. This work was supported by National Institute Health, grant number R01AI162787 from National Institute of Allergy and Infectious Diseases (NIAID). Research reported in this publication was supported by the University of Michigan Advanced Genomics Core, the UM Single Cell Spatial Analysis Program and the National Cancer Institutes of Health under Award Number P30CA046592 by the use of the following Cancer Center Shared Resource: Single Cell and Spatial Analysis Shared Resource.

## Conflict of interest statement

The authors declare no conflicts of interest.

## Notes

### Competing Interest Statement

The authors have declared no competing interest.

